# JQ1 reduces Epstein-Barr virus-associated lymphoproliferative disease in mice without sustained oncogene repression

**DOI:** 10.1101/152306

**Authors:** Amanda He, JJ L. Miranda

**Affiliations:** Department of Cellular and Molecular Pharmacology, University of California, San Francisco, CA, USA; Gladstone Institute of Virology and Immunology, San Francisco, CA, USA

**Keywords:** JQ1, BET inhibitor, Epstein-Barr virus, lymphoproliferative disease, oncogene

## Abstract

Small molecule inhibitors of bromodomain and extra-terminal (BET) proteins are seeing increased investigation in clinical trials for treatment of hematological malignancies. These compounds also repress oncogene expression driven by the human Epstein-Barr virus (EBV) in cell culture. We therefore tested the efficacy of the prototypical BET inhibitor JQ1 against a mouse xenograft model of post-transplantation lymphoproliferative disorder. JQ1 potently inhibits growth of lymphoblastoid cell lines (LCLs) in culture at low nM concentrations. Growth of other cell lines with similar EBV type III latency transcription programs is comparably inhibited. JQ1 also slows tumor development of an LCL xenograft in immunocompromised mice, but oncogene repression is not observed in endpoint biopsies. We find reduction of EBV-associated lymphoproliferative disease in an animal model encouraging of further studies.

---

Emerging preclinical evidence highlights the potential for inhibitors of BET proteins in treating hematological malignancies. These compounds occlude the ligand-binding pockets of bromodomains and prevent recruitment toward acetylated lysine substrates [1, 2]. This activity is leveraged in therapeutic approaches to repress oncogene expression by interfering with transcriptional enhancers. For example, tumor growth is inhibited in mouse models of lymphoma dependent on overexpression of the *MYC* oncogene driven by the BET protein BRD4 [3, 4, 5]. Whether BET inhibitors can similarly repress induced oncogenes to yield efficacy against EBV-associated lymphoproliferative disease *in vivo* remained to be tested.

Efficacy trials of BET inhibitors with EBV-associated malignancies in animal models have so far been limited. EBV immortalizes B cells and is associated with a proportion of malignancies and lymphoproliferative diseases. To facilitate cell growth, EBV increases expression of *MYC* and other oncogenes by producing transcription factors that bind super-enhancers containing BRD4 [6]. JQ1 displays antiproliferative effects against the EBV-infected Raji cell line in a mouse xenograft model of Burkitt lymphoma [3]. Raji viral genomes, however, contain a deletion that results in an atypical transcription profile [7] not commonly found in tumors. LCLs, B cells immortalized by EBV *in vitro*, display a viral transcription program similar to that observed in lymphoproliferative disease among immunosuppressed individuals [8] and serve as commonly tested mouse xenograft models for such lesions [9]. The GM12878 LCL in particular has been heavily examined as part of ENCODE consortium [10]. In this study, we test JQ1 for efficacy in this mouse model of post-transplantation lymphoproliferative disorder.

JQ1 inhibits growth of LCLs at low nM concentration in cell culture. Treatment with JQ1 reduces GM12878 cell growth at a concentration of 500 nM in cell culture [6]. We determined a complete dose response against the active (+)-JQ1 isomer to define a more quantitative measure of potency. Cells at a density of 0.3×10^6^/mL were treated with JQ1 (Selleck Chemicals) in DMSO and counted with a hemacytometer after 3 days. Changes in cell density were normalized against the DMSO vehicle control. We measured an IC_50_ of 134 ± 26 nM (Figure 1(A)). Similar results were obtained with the 721 LCL (data not shown). Low nM IC_50_ concentrations suggest potential potency for use *in vivo*.

**Figure 1.**
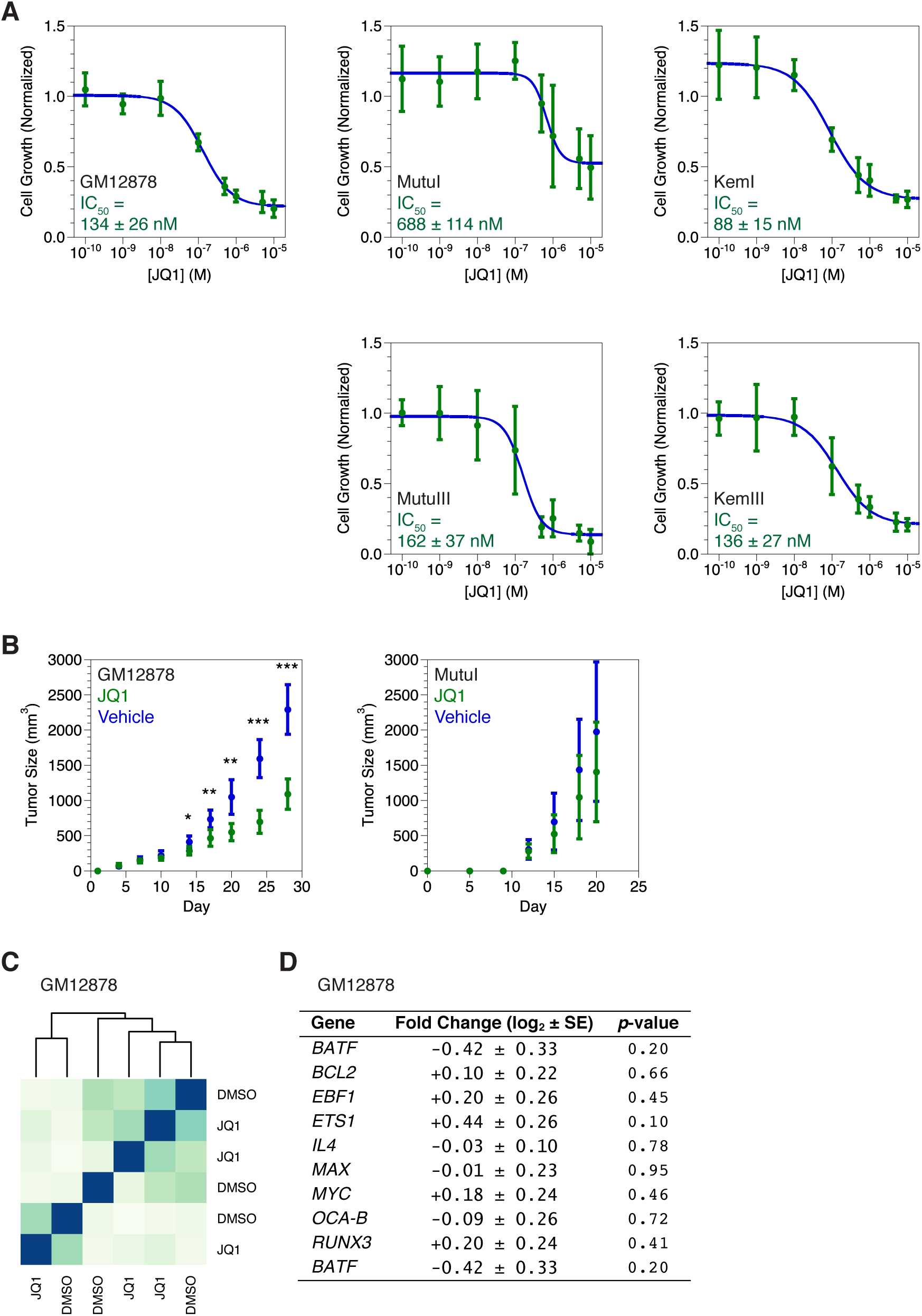
JQ1 reduces growth of EBV-infected LCLs in cell culture and mice. (A) Growth of the GM12878, MutuI, MutuIII, KemI, and KemIII lines in cell culture after 3 days of JQ1 treatment. Expansion is expressed as cell density increase normalized to the vehicle control. Error bars represent the standard deviation of n = 4 replicates. (B) Growth of engrafted GM12878 and MutuI cells in NSG mice treated with JQ1. Expansion is measured as tumor volume. Error bars represent the standard deviation of n = 8 mice. *, **, and *** indicate *p*-values < 0.05, 0.005, and 0.0005, respectively. (C) Hierarchal clustering of RNA-seq transcriptomes in GM12878 tumors from mice treated with JQ1. Heatmap depicts sample-to-sample distances with darker blue indicating greater similarity. (D) Transcriptional changes in GM12878 tumors from mice treated with JQ1 as measured by RNA-seq. Listed are genes important for LCL growth and survival associated with EBV super-enhancers in GM12878 cells. Fold change is expressed as the difference upon JQ1 treatment. Standard error of fold change represents the standard deviation of n = 3 replicates. Reported *p*-values are not adjusted for multiple hypothesis testing.

We further queried the specificity of JQ1 against lymphoma cell lines harboring EBV to determine the generality of responsiveness. Variant strains express different stereotyped transcription programs referred to as latency types. Type III latency expresses transcription factors that assemble EBV super-enhancers, which are sensitive to JQ1 treatment, to possibly promote oncogene expression. Type I latency does not express the same factors. We examined two sets of isogenic Burkitt lymphoma lines that exhibit different viral transcription programs [7]: MutuI paired with MutuIII and KemI paired with KemIII. In all four lines, JQ1 slows growth with IC_50_ values in the nM range (Figure 1(A)). We expected higher IC_50_ values with type I latency lines. This held true for the Mutu pair, 688 ± 114 vs. 162 ± 37 nM, but not the Kem pair, 88 ± 15 vs. 136 ± 37 nM. EBV-mediated regulation of oncogene super-enhancers may therefore be situational. Upon infection to generate LCLs, when oncogene enhancers are not previously active, EBV proteins transform these regulatory elements into super-enhancers for immortalization. In Burkitt lymphoma lines, where oncogene super-enhancers could already be active, EBV proteins may not create additive effects with preexisting mechanisms of oncogenesis. The Kem lines possibly do not display a large difference between latency types because already active super-enhancers reduce the effect of EBV transcription factors that Mutu lines may leverage.

Even though type III latency lines do not consistently yield greater susceptibility to JQ1 compared to isogenic type I lines, all type III lines we examined reveal potent growth inhibition with IC_50_ values of ~100 nM. Our results are consistent with previously observed efficacy against the Daudi and Raji cell lines [6] that express atypical transcription programs [7] consisting of some but not all EBNA proteins that bind EBV super-enhancers. Type III latency, identifiable in patient samples, may provide predictive power for BET inhibitor sensitivity in the personalized medicine of B cell lymphoma treatments.

JQ1 decreases the growth of EBV-associated lymphoproliferative disease in a mouse xenograft model. We implanted 5×10^6^ GM12878 cells via subcutaneous injection into one flank of 7–9-week-old NSG (NOD.Cg-*Prkdc^scid^ Il2rg^tm1Wjl^*/SzJ) immunocompromised mice. JQ1 in 10% (2-hydroxypropyl)-β-cyclodextrin (Sigma-Aldrich) was administered at a dose of 50 mg/kg via intraperitoneal injection every day and tumor volumes measured with calipers. Lymphoproliferative disease was compared to a vehicle-treated cohort. JQ1 reduces tumor development significantly after ~2 weeks as determined by Student’s *t*-test (Figure 1(B)). Tumor size decreases ~2-fold after 3–4 weeks. We obtained contrasting results with a xenograft model of Burkitt lymphoma. When MutuI cells were implanted into mice with the same protocol, we observed no statistically significant difference in tumor size with JQ1 treatment (Figure 1(B)). The higher IC50 value in cell culture may explain the lack of efficacy against MutuI cells in xenografts. For the GM12878 LCL treated with JQ1, the low nM IC_50_ value *in vitro* translates to reduction of EBV-associated lymphoproliferation *in vivo*.

Transcriptional profiling of tumors surprisingly detected minimal perturbation after JQ1 treatment. We searched for potential antitumor mechanisms in the GM12878 xenograft with RNA deep sequencing (RNA-seq). Specimens were excised in ~30 mm^3^ slices from mice after 1 month of JQ1 treatment and ~5 hours after the final dose. Biopsies were stored in RNA*later* (Qiagen) at -80 °C and disrupted with a pellet pestle (Kimble Chase) in RLT Buffer (Qiagen). RNA was purified, processed, and sequenced according to our RNA-seq protocol [7]. Data analysis was performed with the <u>usegalaxy.org</u> instance of Galaxy [11]: reads were aligned to the UCSC hg19 human genome reference sequence with HiSAT version 2.0.3 [12], gene-level expression abundance measured using the UCSC hg19 RefSeq annotation with featureCounts version 1.4.6-p5 [13], and differentially expressed genes identified with DESeq2 version 1.14.1 [14]. Every data set yielded ~10–40 million mapped reads. Comparing three samples each of JQ1- and vehicle-treated biopsies, hierarchal clustering reveals little distinction between transcriptome profiles based on JQ1 treatment (Figure 1(C)). Concordant with this measure of similarity, we detected only 22 differentially expressed genes with a log2 fold change > 1 and *p*-value adjusted for multiple testing < 0.05. JQ1 decreases transcript levels of F5, NEURL3, ANXA3, GATM, KCNJ2, ARBB1, PRRX1, MAP7D2, NOL4, IL1R2, POU2F3, KCNMB1, ANO10, BARX2, ATP1B1, NHS, SVIL, PARD3, TUBB6, and CBLN2; JQ1 increases transcript levels of ZAK and PIP5K1B. Some of these genes have been implicated in other types of cancer and may be suitable for follow-up experiments investigating roles in LCL growth. The minute number of differentially regulated genes indicates that JQ1 causes only small changes in transcription as measured with RNA-seq.

JQ1 does not repress transcription of oncogenes associated with EBV super-enhancers in endpoint biopsies. We individually examined the effect of JQ1 on several genes associated with LCL growth (Figure 1(D)). *MYC* expression levels are not reduced. We also examined *BCL2* and *RUNX3*, genes also controlled by EBV super-enhancers [6]. Again, we found no repression. Even expanding our search to genes important for LCL growth proximal to EBV super-enhancers [6] did not detect any significant transcriptional changes. JQ1 pharmacokinetics may be responsible for the lack of transcriptional perturbation, but *MYC* repression occurs in physiologically similar Raji xenografts of NOD/SCID mice 4 and 8 hours after dosing with 25 mg/kg JQ1 [3]. We favor the possibility that the GM12878 LCL escapes oncogene repression through rapidly acquired resistance to JQ1 treatment. Our observation that GM12878 xenografts do not display sustained oncogene repression should emphasize this weakness as a point of optimization in BET inhibitor therapeutic strategies.

Although our experiments demonstrate only partial reduction of tumor growth in a mouse xenograft model, we still advocate further research into the use of BET inhibitors for treating EBV-associated post-transplantation lymphoproliferative disorder. BET inhibitors are already seeing increasing use in clinical trials, generating a wealth of data in the context of B cell malignancies. Moreover, not only do BET inhibitors target oncogene expression driven by EBV proteins [6], but our own work shows that the same compounds also block EBV lytic gene expression [15], a contributing factor [9] to lymphoproliferative disease. This potential therapeutic deserves investigation in further preclinical animal models.

## Acknowledgments

We thank Byron C. Hann and the Preclinical Therapeutics Core at UCSF for help with designing, conducting, and discussing *in vivo* studies. We are grateful to the UCSF Center for Advanced Technology and the Gladstone Genomics Core for use of shared equipment. This project was funded by the UCSF Program for Breakthrough Biomedical Research, funded in part by the Sandler Foundation. Our work was supported by Grant #IRG-97-150-13 from the American Cancer Society.

